# Immuno-EM characterization of vasopressin neurons in macaques by use of formaldehyde-fixed tissues stored for several years

**DOI:** 10.1101/2021.06.07.447309

**Authors:** Akito Otubo, Sho Maejima, Takumi Oti, Keita Satoh, Yasumasa Ueda, John F. Morris, Tatsuya Sakamoto, Hirotaka Sakamoto

## Abstract

Translational research often requires the testing of experimental therapies in primates, but research in non-human primates is now stringently controlled by law around the world. Tissues fixed in formaldehyde without glutaraldehyde have been thought to be inappropriate for use in electron microscopic analysis, particularly those of the brain. Here we report the immunoelectron microscopic characterization of arginine vasopressin (AVP)-producing neurons in macaque hypothalamo-pituitary axis tissues fixed with 4% formaldehyde and stored at –25°C for several years. The size difference of dense-cored vesicles between magnocellular and parvocellular AVP neurons was detectable in their cell bodies and perivascular nerve endings located, respectively, in the posterior pituitary and median eminence. Furthermore, glutamate and the vesicular glutamate transporter 2 were colocalized with AVP in perivascular nerve endings of both the posterior pituitary and the external layer of the median eminence, suggesting that both magnocellular and parvocellular AVP neurons are glutamatergic in primates. Both ultrastructure and immunoreactivity can therefore be sufficiently preserved in macaque brain tissues stored long-term for light microscopy. Taken together, these results suggest that this methodology could be applied to the human post-mortem brain and be very useful in translational research.

## 1. Introduction

Studies using macaque monkeys as model non-human primates are essential because of their high applicability to human clinical research which often requires the testing of experimental therapies in primates. However, primate research is now stringently controlled by law around the world (Abbott, 2014). Rodents, in which research is less restricted, are often not analogous enough to humans in their physiology to allow an extrapolation from rodent to human. Medical science is urged to make minimal use of animals such as primates, and basic studies on non-human primates appear to be permitted only if the work could not be carried out in any other species. Thus, although tissues from primates are often essential to achieve relevant results, there are, nevertheless, considerable problems in their use.

Arginine vasopressin (AVP), an anti-diuretic hormone, is released, not only into the blood stream, but also into the central nervous system where, in mammals, it has been shown to be important for stress coping, aggression, courtship behavior, learning, bonding, and various socio-sexual behaviors (Walum & Young, 2018). AVP is produced mainly by magnocellular neurosecretory neurons in the supraoptic nucleus (SON) and paraventricular nucleus (PVN) of the hypothalamus (Buijs et al., 1983). The *AVP* gene encodes a precursor containing AVP, AVP-associated neurophysin II (NPII), and a glycopeptide copeptin (Breslow, 1993; Davies et al., 2003; Land et al., 1982). The expression and release of AVP by magnocellular neurosecretory neurons in the SON and PVN are regulated by physiological conditions, including plasma osmotic pressure and blood pressure (Koshimizu et al., 2012). The magnocellular axons project primarily *via* the internal layer of the median eminence to the posterior pituitary where they release AVP into the systemic circulation. In addition, some parvocellular neurons in the PVN produce AVP and project into extrahypothalamic areas where the AVP and/or other co-packaged molecules regulate brain function as neuromodulators (Morris, 2020).

Corticotrophin-releasing factor (CRF) is a strong stimulator of adrenocorticotrophic hormone (ACTH) secretion from the anterior pituitary when released onto portal capillaries in the median eminence in response to the stress (Vale et al., 1981). A population of parvocellular neurons in the anteromedial part of the PVN produces CRF and may, depending on the extent of stress, also produce AVP. Both peptides released into the hypothalamo-hypophysial portal circulation play an important synergistic role in stress resilience (Fellmann et al., 1984; Itoi et al., 2014; Mouri et al., 1993; Whitnall et al., 1985). Because intense AVP- and CRF-immunoreactivity have been observed in the external layer of the macaque median eminence, the peptides are probably co-released into the portal circulation to amplify ACTH release from the primate anterior pituitary (Otubo et al., 2020; Zimmerman et al., 1973).

In rodents, the presence of glutamate-immunoreactivity in magnocellular neuroendocrine cells of the SON suggests that AVP neurons also produce glutamate as a neurotransmitter (Meeker et al., 1991; Meeker et al., 1989). Within the neurosecretory endings of the posterior pituitary, glutamate immunoreactivity is specifically localized to electron-lucent microvesicles with no overlap onto the dense-cored neurosecretory vesicle (dcv) population in rats (Meeker et al., 1991). Immunocytochemical co-localization of CRF and the vesicular glutamate transporter 2 (VGLUT2) in rats suggests that the co-release of CRF and glutamate may function to regulate postsynaptic targets (Valentino et al., 2001). It is currently unclear whether glutamate has a similar or other functions in the primate AVP/CRF system.

Tissues fixed in formaldehyde without glutaraldehyde have long been thought to be inappropriate for use in electron microscopic analysis, especially in the brain (Maunsbach, 1966a, 1966b). Here we report the immunoelectron microscopic characterization of AVP-producing neurons in the primate hypothalamo-pituitary axis tissue fixed with formaldehyde and stored at –25°C for several years. Special attention was paid to the size of dcv in AVP- producing magno- and parvocellular neurons and to the colocalization of CRF with AVP- related gene products in the dcv. We show that immunoelectron microscopy of formaldehyde-fixed tissue can confirm the size difference in dcv between magno- and parvocellular AVP neurons in Japanese macaque monkeys. Furthermore, we show that, in macaque monkeys, AVP/CRF and the VGLUT2/glutamate are co-localized in both the magnocellular endings of the posterior pituitary and the parvocellular endings in the external layer of the median eminence.

## 2. Results

### 2.1. Antibody characterization and the expression of VGLUT2 at the protein level in the posterior pituitary

Full details of all the antibodies used in this study are shown in Table 1. We first validated by Western blot analysis the specificity for the VGLUT2 and NPII antibodies in Japanese macaque monkeys. Western blot analysis demonstrated the specificity for the guinea pig polyclonal antibody against VGLUT2 and the expression of VGLUT2 at the protein level in the posterior pituitary; a single strong immunoreactive band was detected at the expected molecular weight of ~56 kDa (Balaram et al., 2013) (Fig. 1). The mouse monoclonal antibody against AVP-neurophysin II (NPII) also detected a single major band at the expected molecular weight of ~12 kDa on the blots of posterior pituitary (Kawakami et al., 2021; Otubo et al., 2020) (Fig. 1).

**Table 1.**
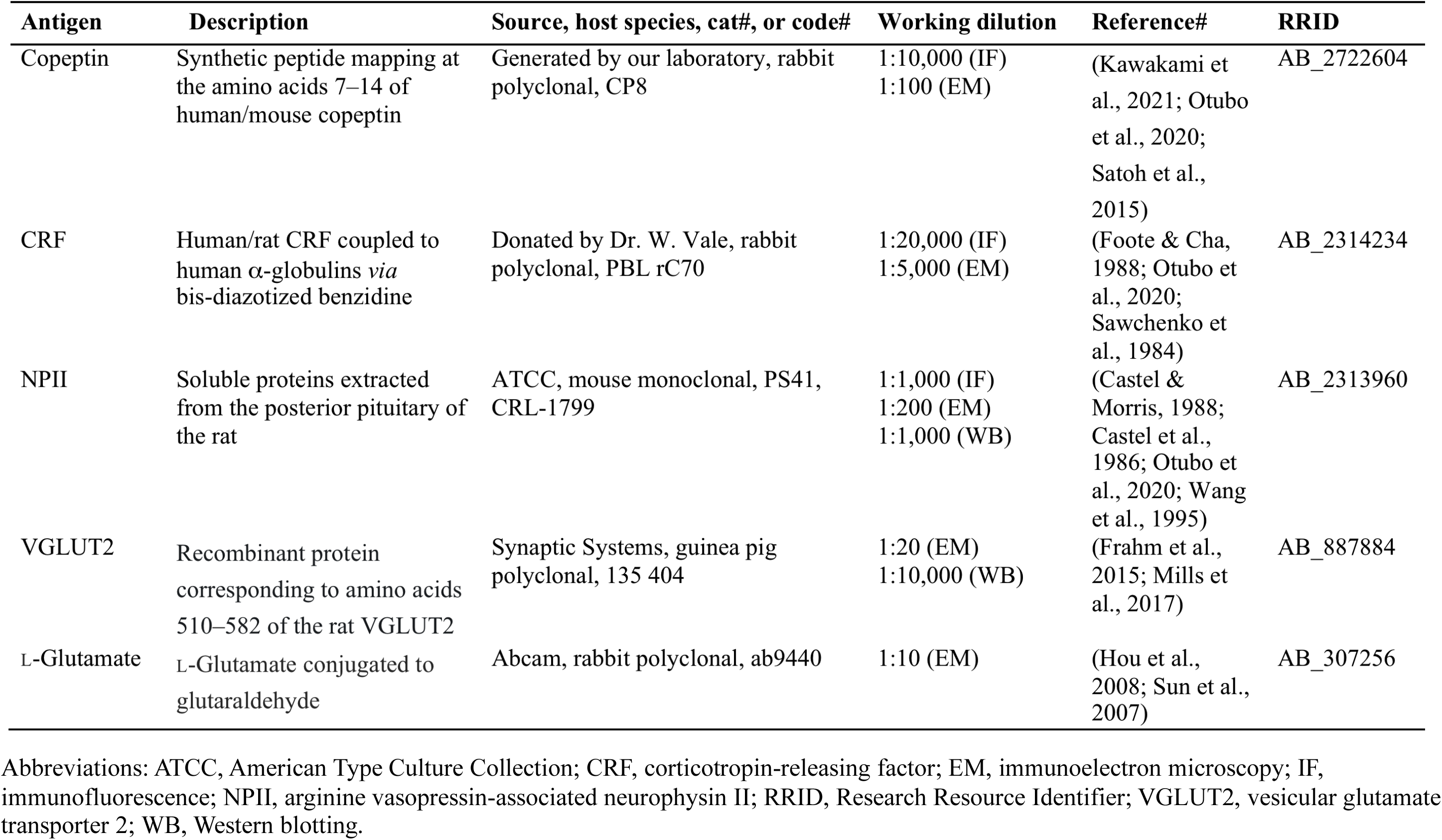
Primary antibodies used in this study.

**Figure 1.**
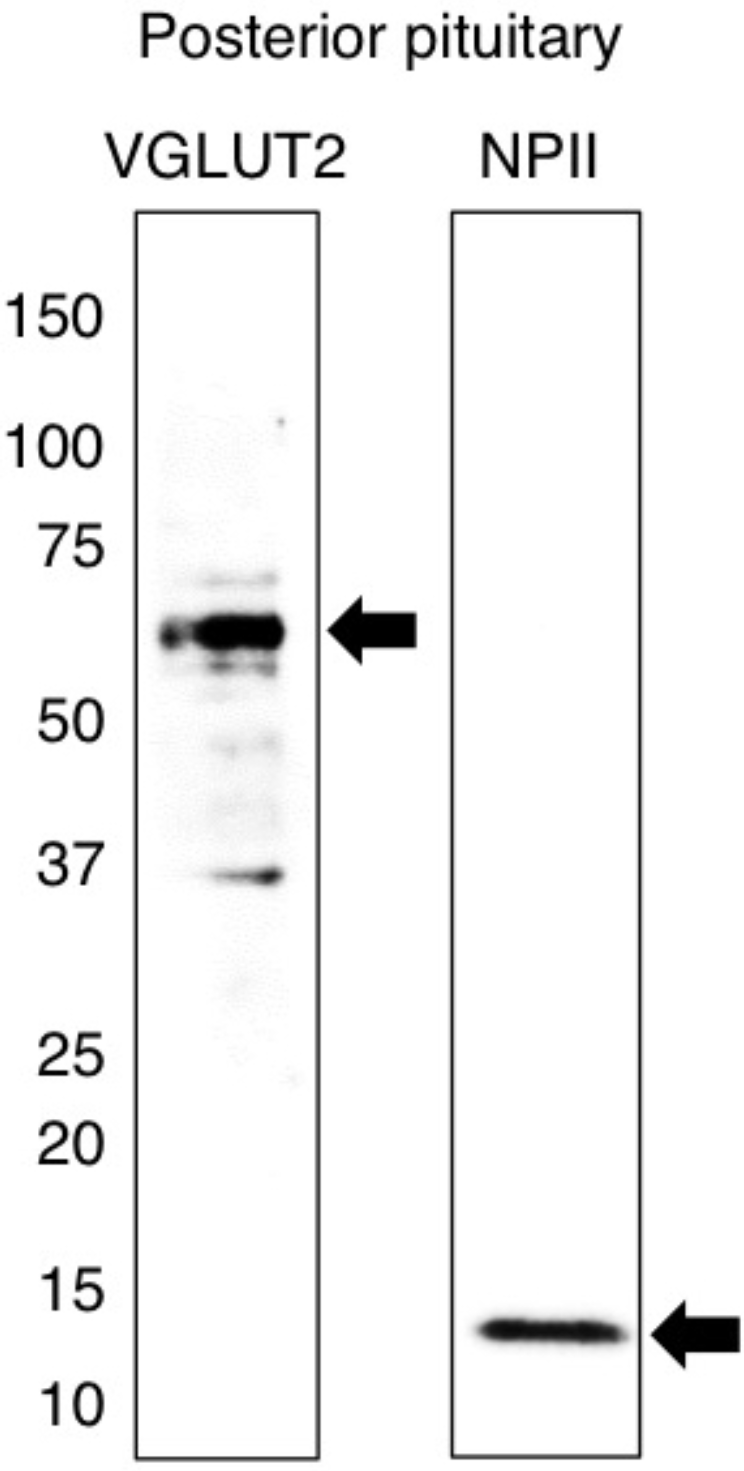
Western immunoblotting of vesicular glutamate transporter 2 (VGLUT2) and vasopressin-associated neurophysin (NPII). The number on the left indicates the molecular weight (kDa). Extracts of protein from the posterior pituitary of the Japanese macaque monkey were transferred onto polyvinylidene difluoride membranes and probed with the guinea pig polyclonal antiserum against VGLUT2 or with the mouse monoclonal antibody against NPII. The antisera recognized a single major band at the expected molecular weight of VGLUT2 (~56 kDa) or NPII (~12 kDa) on a Western blot of the macaque posterior pituitary.

### 2.2. Anatomy of AVP-producing neurons in the macaque hypothalamus

Immunostaining for NPII was performed to study the localization of AVP neurons in the hypothalamus of Japanese macaque monkeys. Numerous magnocellular (large-sized) AVP neurons were observed in the PVN (Fig. 2A) and SON (Fig. 2B). Parvocellular (small-sized) AVP neurons were also observed in the PVN (Fig. 2C). The median eminence can be divided into an internal and an external layer. The internal layer is the zone through which the axons of magnocellular neurons densely project into the posterior pituitary (Fig. 2D) (Buijs et al., 1983; Morris, 2020; Zimmerman et al., 1973). The external layer of the median eminence is where the axons of many parvocellular neurosecretory neurons, including parvocellular AVP/CRF neurons of the PVN, end on hypothalamo-hypophysial portal vessels and release anterior pituitary-hormone-regulating hormones (Fig. 2D) (Buijs et al., 1983; Morris, 2020; Zimmerman et al., 1973).

**Figure 2.**
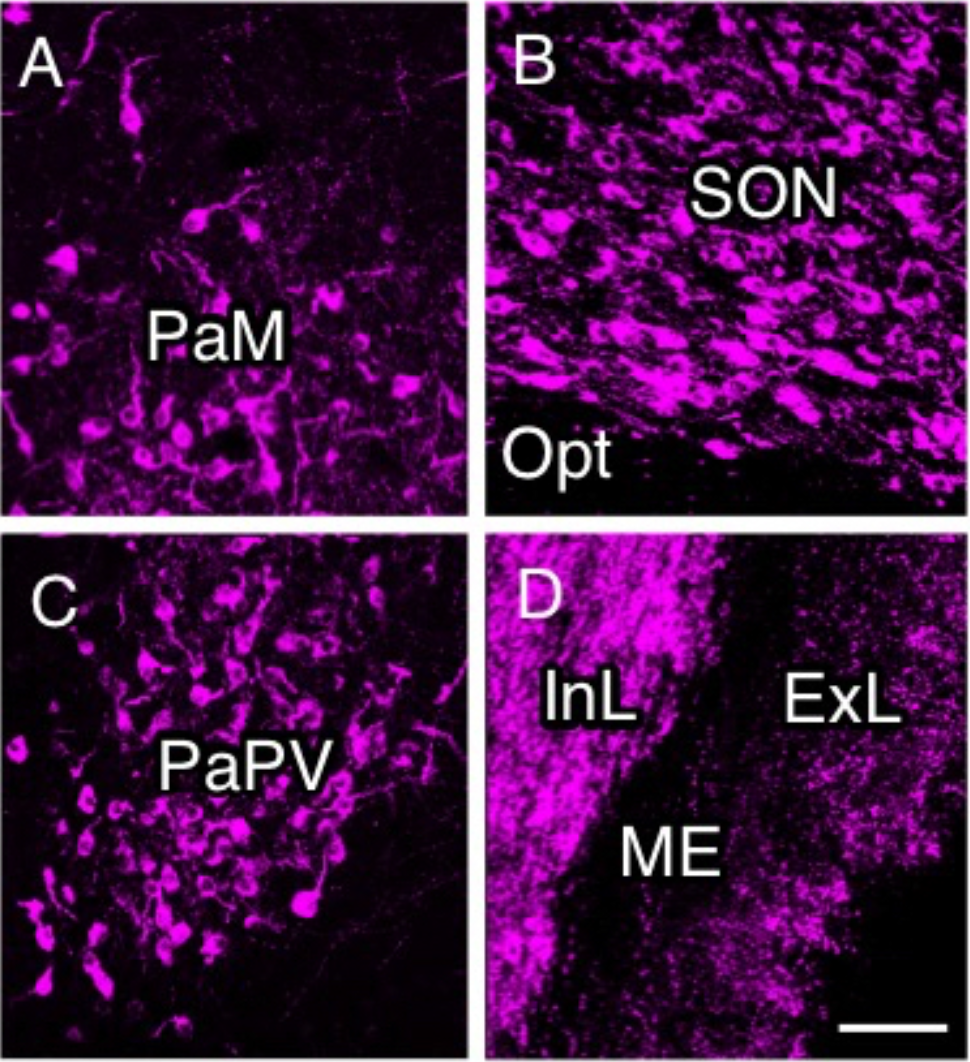
Immunofluorescence for vasopressin-associated neurophysin (NPII) in the macaque hypothalamus. NPII-immunoreactivity was observed in the paraventricular nucleus (PVN) of the hypothalamus; PaM, magnocellular part of the PVN (A); supraoptic nucleus (B); PaPV, parvocellular part of the PVN, ventral division (C); and median eminence (ME) (D). Opt, optic nerve; InL, internal layer of the ME; ExL, external layer of the ME. *Scale bar*, 100 μm.

Immunoelectron microscopy of the posterior pituitary showed many AVP dcv containing both NPII (PS41)- and copeptin (CP8)-immunoreactivity (Fig. 3A, B). In other neighbouring endings, no NPII- or copeptin-immunoreactivity was detectable (Fig. 3C). In the median eminence, many dcv showing PS41/CP8 double-immunoreactivity were observed in both the internal and external layers (Fig. 4). However, the AVP dcv in the internal layer (magnocellular) of the median eminence (Fig. 4B) were larger than those in the external layer (parvocellular) of the median eminence (Fig. 4D).

**Figure 3.**
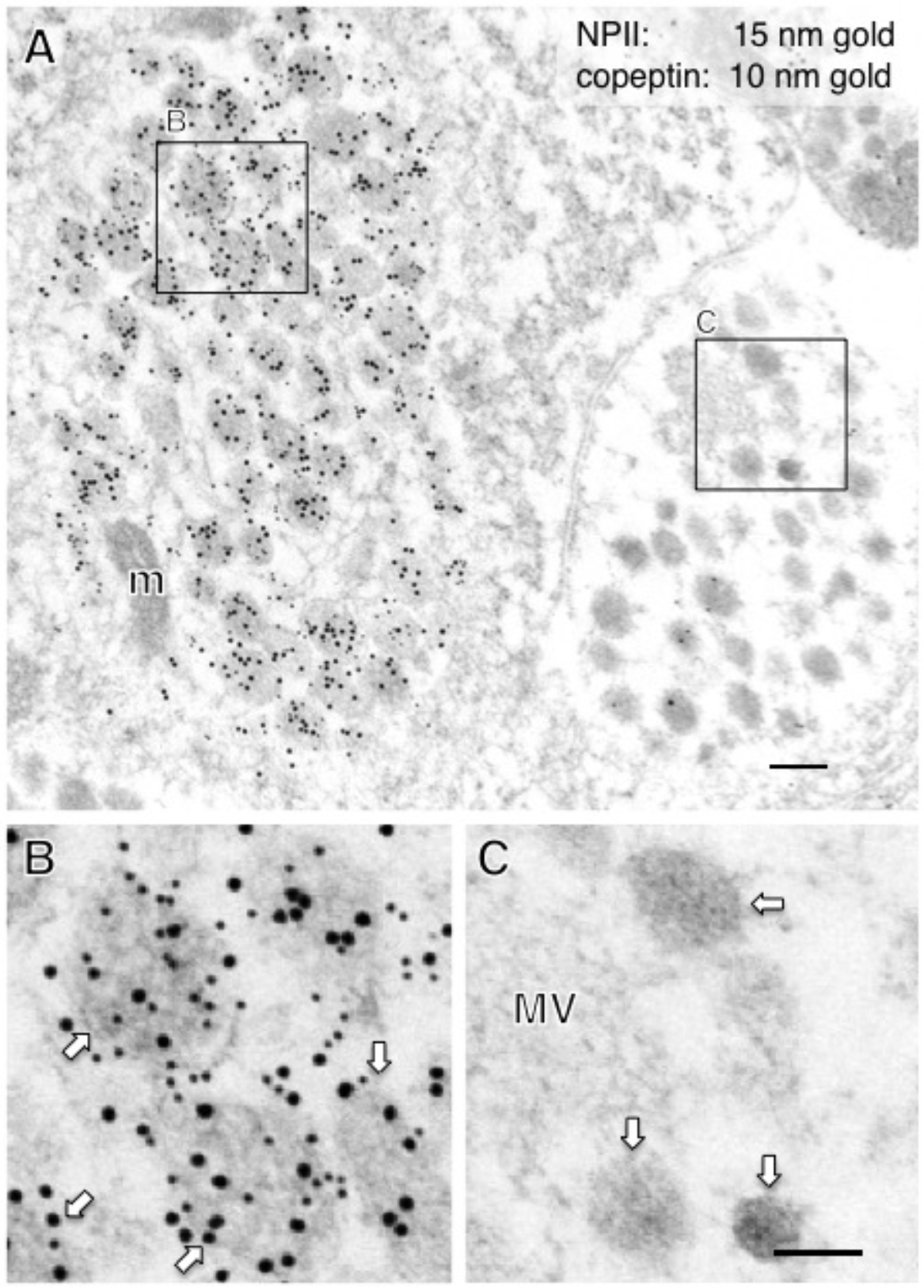
Double-label immunoelectron microscopy for vasopressin-associated neurophysin (NPII) and copeptin in the macaque posterior pituitary. Numerous neurosecretory vesicles located in some of the varicosities were doubly immunopositive for copeptin (10-nm gold particles) and NPII (15-nm gold particles). The outlined areas in (A) are enlarged in (B) and (C). In contrast, in other presumably oxytocin-containing varicosities, no NPII/copeptin-immunoreactivity was detected (A and C). *Scale bars*, 200 nm, and 100 nm in enlarged images. *Arrows* indicate dense-cored neurosecretory vesicles. m, mitochondrion; MV, clustered microvesicles.

**Figure 4.**
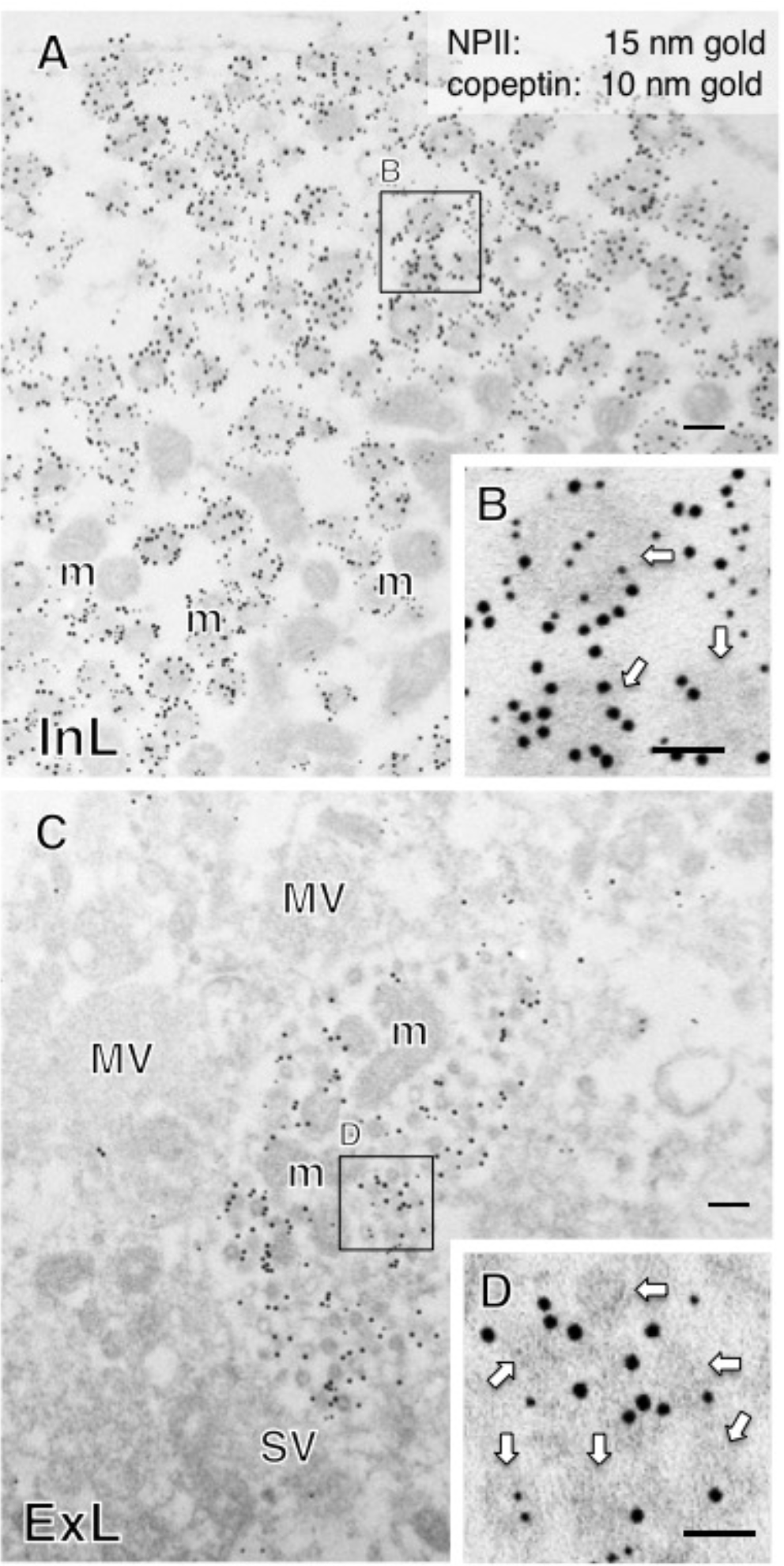
Double-label immunoelectron microscopy for vasopressin-associated neurophysin (NPII) and copeptin in the internal layer (InL; A, B) and external layer (ExL; C, D) of the macaque median eminence. Numerous dense-cored neurosecretory vesicles located in some of the varicosities were immunopositive for both copeptin (10-nm gold particles) and NPII (15-nm gold particles). The outlined area in (A) and (C) is enlarged in (B) and (D), respectively. *Scale bars*, 200 nm, and 100 nm in enlarged images. *Arrows* indicate dense-cored neurosecretory vesicles. m, mitochondrion; MV, clustered microvesicles.

### 2.3. Size characterization of dcv

In this study, immunoelectron microscopic characterization of AVP-producing neurons was performed in macaque hypothalamo-pituitary axis tissues fixed with 4% formaldehyde and stored at –25°C for several years. We found that both ultrastructure and immunoreactivity are sufficiently preserved in macaque brain tissues stored long-term for light microscopy. Approximately 200 dcv in the neurosecretory axons in the posterior pituitary (magnocellular), the internal layer of the median eminence (magnocellular), and the external layer of the median eminence (parvocellular) were selected and their major axis diameters were measured. The average of major axis was 196 ± 2 nm in the posterior pituitary, 197 ± 2 nm in the internal layer of the median eminence, but significantly smaller (93 ± 1 nm) in the external layer of the median eminence (Fig. 5; Table 2).

**Table 2.**
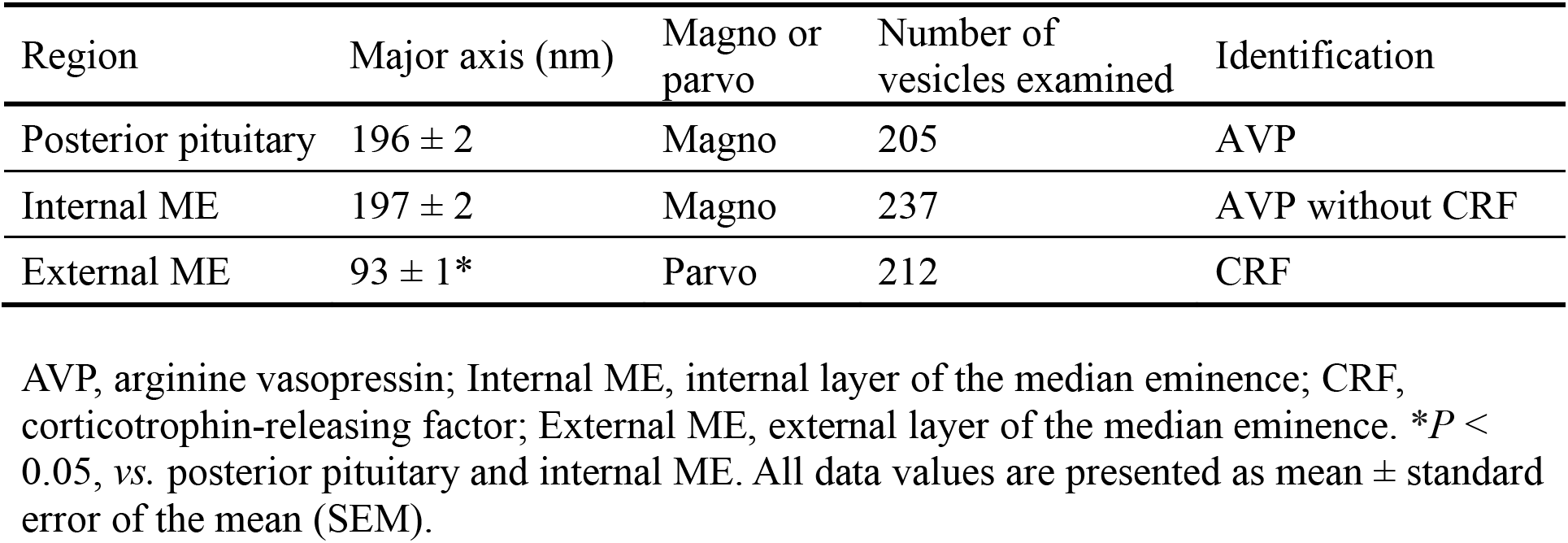
Identification of cell types (magno- or parvocellular neurons) based on the major axis of dense-cored neurosecretory vesicles.

| Region | Major axis (nm) | Magno or parvo | Number of vesicles examined | Identification |
| --- | --- | --- | --- | --- |
| Posterior pituitary | 196 ± 2 | Magno | 205 | AVP |
| Internal ME | 197 ± 2 | Magno | 237 | AVP without CRF |
| External ME | 93 ± 1* | Parvo | 212 | CRF |

- 5 AVP, arginine vasopressin; Internal ME, internal layer of the median eminence; CRF, corticotrophin-releasing factor; External ME, external layer of the median eminence. \* $P < 0.05$ , vs. posterior pituitary and internal ME. All data values are presented as mean ± standard error of the mean (SEM).

**Figure 5.**
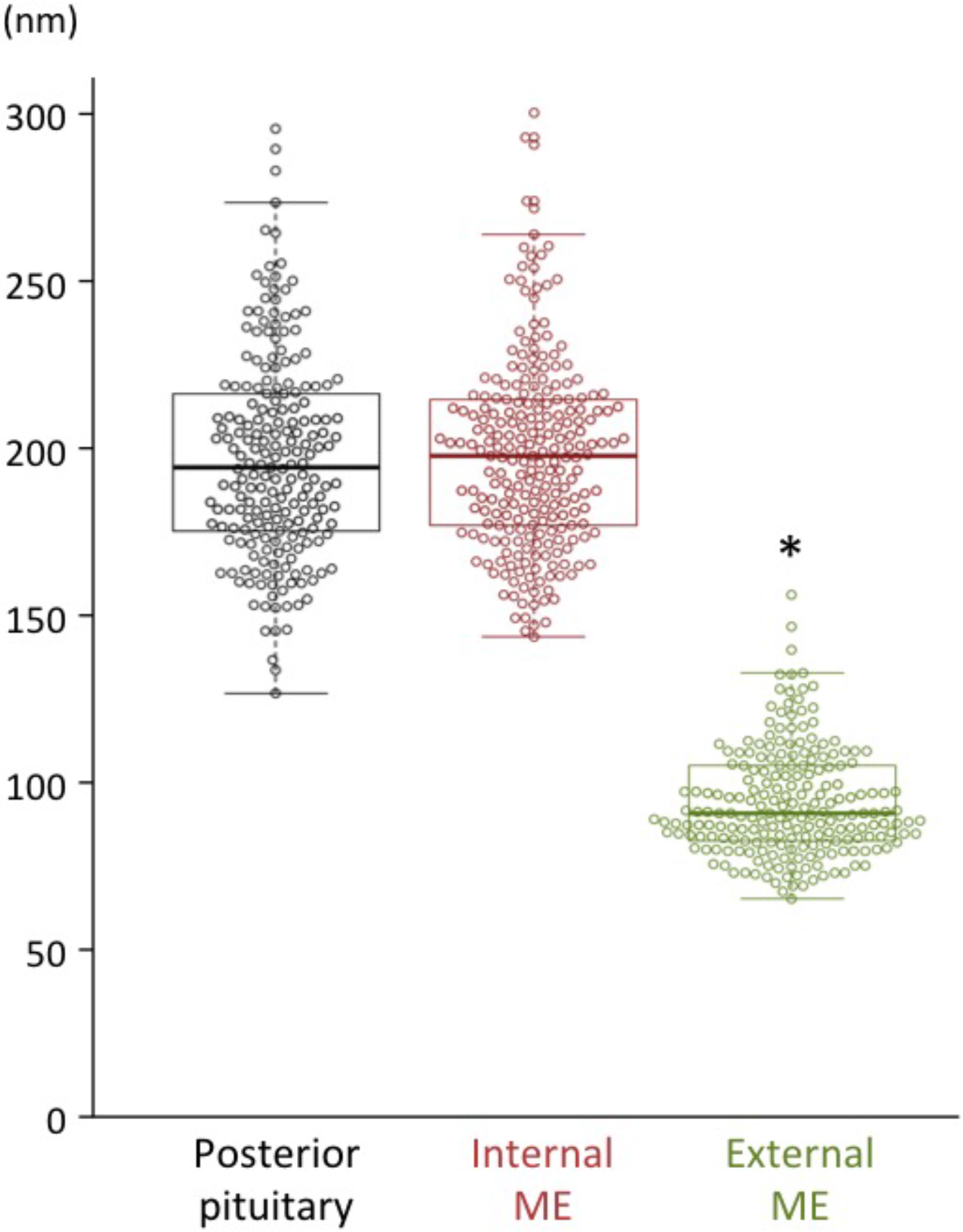
Bee-swarm and box plots displaying the major axis of dense-cored neurosecretory vesicles distributed in the posterior pituitary (black), internal layer of the median eminence (ME) (red), and external layer of the ME (green) in macaque monkeys. The major axis of dense-cored vesicles in the posterior pituitary and the internal layer of the ME (~200 nm) was significantly larger than that of the vesicles in the external layer of the ME (~100 nm). \**P* < 0.05, *vs.* posterior pituitary and internal layer of the ME.

Next, we measured the diameter of dcv in the perikarya of randomly selected SON and PVN AVP neurosecretory neurons (at x5,000 magnification; Table 3). In the SON (7 cell bodies), the average diameter of dcv contained in the cell body was 204 ± 12 nm (11 vesicles), confirming that, as expected, all AVP neurons in the SON are magnocellular (Fig. 6; Table 3). The 18 AVP neurons in the PVN selected for analysis formed two distinct groups: those with larger dcv (180 ± 19 nm; 11 vesicles; Fig. 7A, B) identified as magnocellular; and those with smaller dcv (102 ± 10 nm; 14 vesicles; Fig. 7C, D) identified as parvocellular neurons.

**Table 3.** Characterization of cell types (magno- or parvocellular neurons) based on the major axis of dense-cored neurosecretory vesicles.

| Cell body | Major axis (nm) | Magno or parvo | Number of vesicles examined | SON or PVN | Identification |
| --- | --- | --- | --- | --- | --- |
| #1 | 187 ± 5* | Magno | 29 | SON | AVP |
| #2 | 175 ± 6* | Magno | 35 | SON | AVP |
| #3 | 189 ± 10* | Magno | 24 | SON | AVP |
| #4 | 181 ± 5* | Magno | 26 | SON | AVP |
| #5 | 171 ± 7* | Magno | 25 | SON | AVP |
| #6 | 195 ± 6* | Magno | 30 | SON | AVP |
| #7 | 205 ± 6* | Magno | 46 | SON | AVP |
| #8 | 188 ± 5* | Magno | 49 | PVN | AVP |
| #9 | 170 ± 5* | Magno | 20 | PVN | AVP |
| #10 | 169 ± 8* | Magno | 22 | PVN | AVP |
| #11 | 172 ± 4* | Magno | 24 | PVN | AVP |
| #12 | 184 ± 5* | Magno | 25 | PVN | AVP |
| #13 | 182 ± 5† | Magno | 21 | PVN | AVP |
| #14 | 103 ± 5† | Parvo | 38 | PVN | AVP |
| #15 | 98 ± 4† | Parvo | 42 | PVN | CRF |
| #16 | 109 ± 4† | Parvo | 51 | PVN | CRF |
| #17 | 93 ± 5† | Parvo | 24 | PVN | CRF |
| #18 | 107 ± 5† | Parvo | 40 | PVN | CRF |
| #19 | 103 ± 4† | Parvo | 35 | PVN | CRF |
| #20 | 110 ± 7† | Parvo | 42 | PVN | CRF |
| #21 | 108 ± 5† | Parvo | 77 | PVN | CRF |
| #22 | 93 ± 4† | Parvo | 81 | PVN | CRF |
| #23 | 95 ± 5† | Parvo | 30 | PVN | CRF |
| #24 | 120 ± 6*† | (Parvo) | 40 | PVN | CRF |
| #25 | 113 ± 3*† | (Parvo) | 89 | PVN | CRF |

- 5 AVP, arginine vasopressin; CRF, corticotrophin-releasing factor; SON, supraoptic nucleus; PVN, paraventricular nucleus. \* $P < 0.05$ , vs. external layer of the median eminence; † $P < 0.05$ , vs. posterior pituitary and internal layer of the median eminence. All data values are presented as mean ± standard error of the mean (SEM).

**Figure 6.**
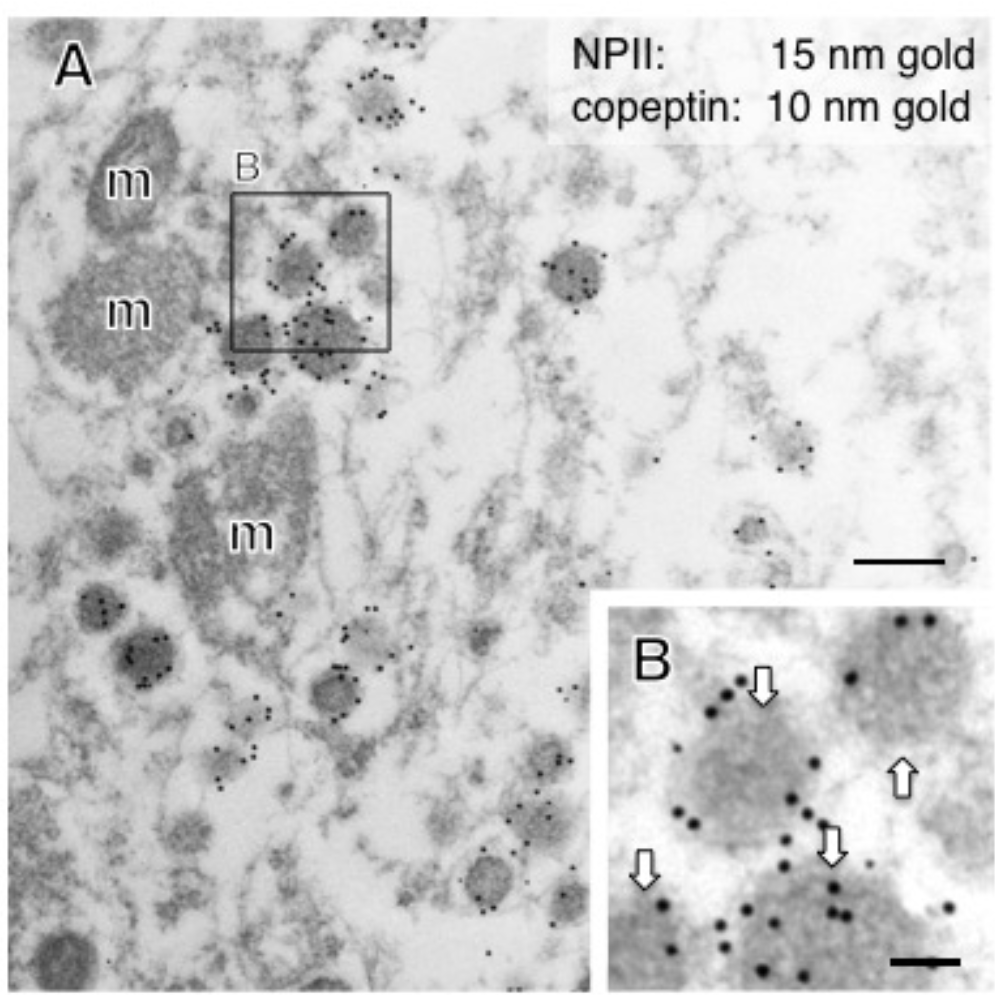
Double-label immunoelectron microscopy for vasopressin-associated neurophysin (NPII) and copeptin in the macaque supraoptic nucleus (SON). Many dense-cored neurosecretory vesicles located in the cell body were doubly immunopositive for copeptin (10-nm gold particles) and NPII (15-nm gold particles). The outlined area in (A) is enlarged in (B). *Scale bars*, 200 nm, and 100 nm in the enlarged image. *Arrows* indicate dense-cored neurosecretory vesicles. m, mitochondrion.

**Figure 7.**
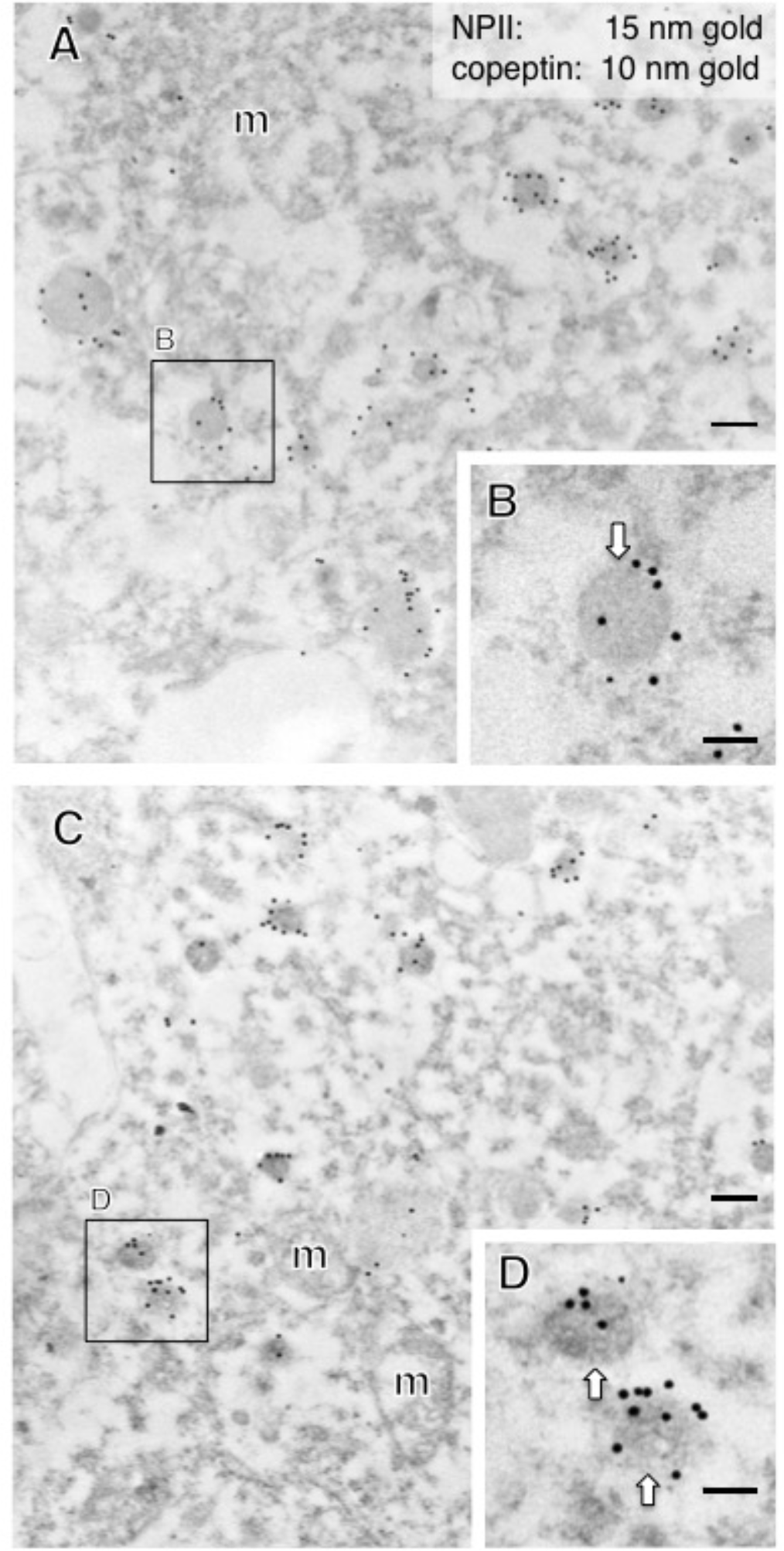
Double-label immunoelectron microscopy for vasopressin-associated neurophysin (NPII) and copeptin in the macaque paraventricular nucleus (PVN) of the hypothalamus. (A) In the cell body of a putative magnocellular vasopressin neuron, many dense-cored neurosecretory vesicles located in the cell body were doubly immunopositive for copeptin (10-nm gold particles) and NPII (15-nm gold particles). (C) Similar results are obtained in the cell body of a putative parvocellular vasopressin neuron. The size of dense-cored vesicles in (A and B) was larger than that of vesicles in (C and D). The outlined areas are enlarged. *Scale bars*, 200 nm, and 100 nm in enlarged images. *Arrows* indicate dense-cored neurosecretory vesicles. m, mitochondrion.

### 2.4. A subpopulation of parvocellular AVP neurons in the PVN co-expresses CRF

In the PVN, of the 18 AVP neuronal cell bodies analyzed, 6 were identified as magnocellular on the basis of vesicle size; none of these showed any CRF-immunoreactivity. Of the 12 PVN perikarya identified as parvocellular on the basis of vesicle size, a subpopulation (especially in the parvocellular part of the PVN) co-expressed CRF (Fig. 8) and such neurons were more intermingled with magnocellular neurons in the macaque monkey than in rodent PVN as reported previously (Foote & Cha, 1988; Kawata & Sano, 1982; Otubo et al., 2020). The terminal regions of parvocellular AVP/CRF neurons in the external layer of the median eminence were next analyzed (Fig. 9). In these cells, the major axis diameter of the sparse dcv was 106 ± 3 nm (16 vesicles) (Fig. 10) for the vesicles in the CRF-immunoreactive terminals, comparable to the 113 ± 9 nm (12 vesicles) in the cell body (Fig. 11). In two putative parvocellular neurons (#24 and #25), the mean major axis of the dcv appeared slightly larger which might relate to the maturity of the vesicles (Table 3) (Cannata & Morris, 1973).

**Figure 8.**
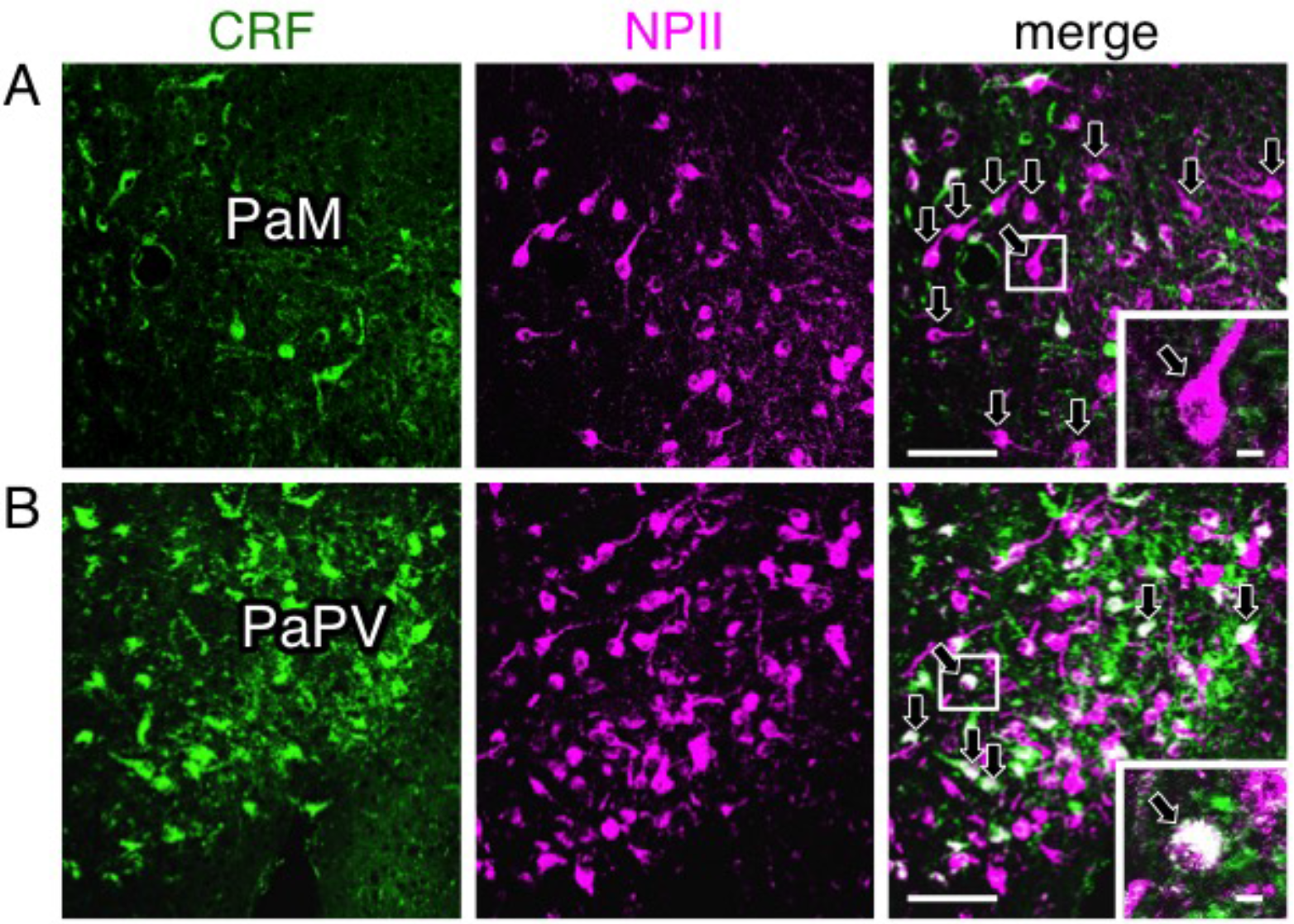
Double-label immunofluorescence for corticotrophin-releasing factor (CRF) and vasopressin-associated neurophysin (NPII) in the paraventricular nucleus (PVN) of the macaque hypothalamus; PaM, magnocellular part of the PVN (A); PaPV, parvocellular part of the PVN, ventral division (B). Immunoreactivity against CRF (green) and NPII (magenta) is merged in each right-hand *panel* (overlap; white). The outlined areas in merged images are enlarged. *Arrows* in (A; merged) indicate perikarya single-immunopositive for NPII in the PaM. *Arrows* in (B; merged) indicate doubly immunopositive for CRF and NPII in the PaPV. *Scale bars*, 100 μm, and 10 μm in enlarged images.

**Figure 9.**
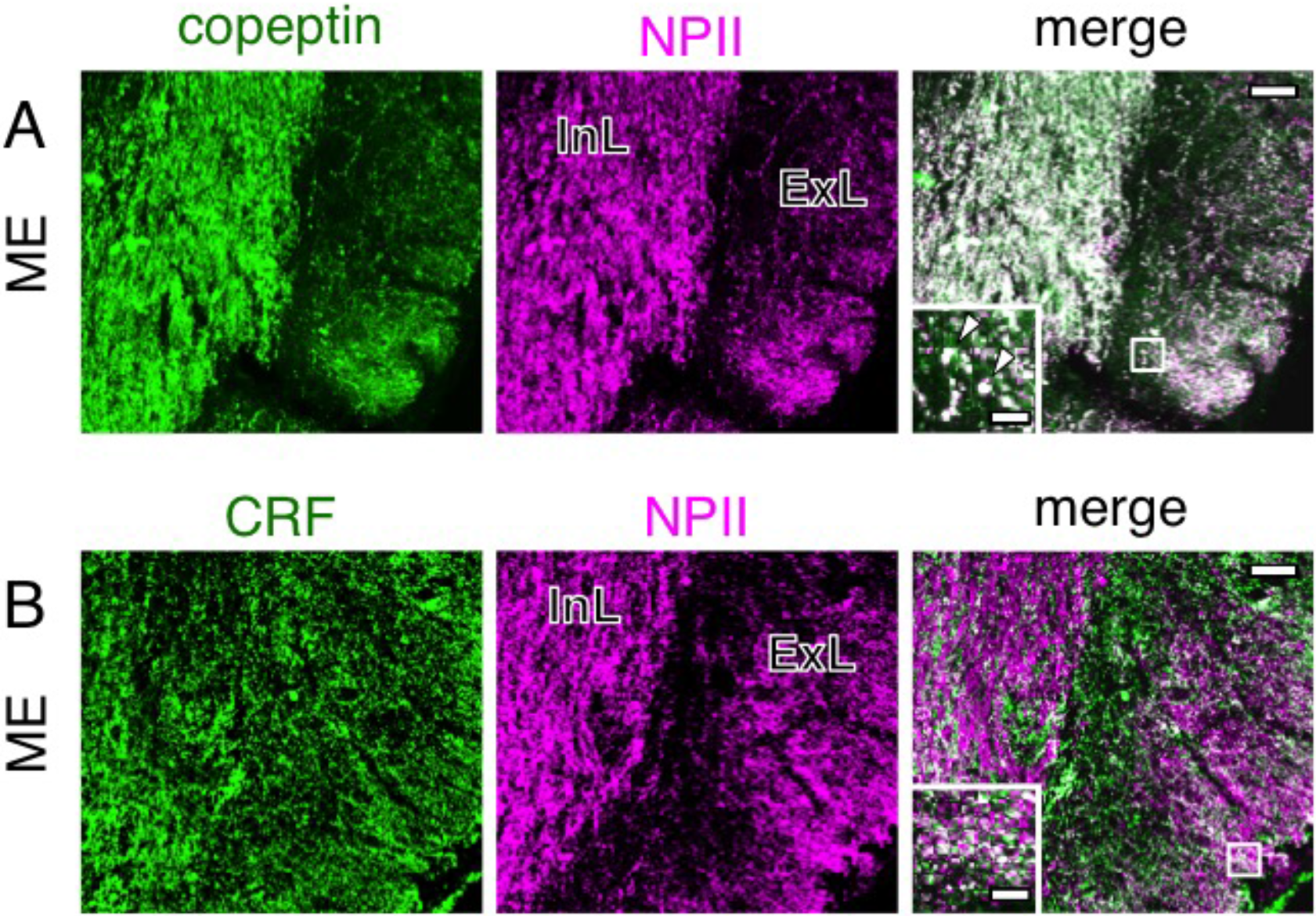
Double-label immunofluorescence for copeptin (A) or corticotrophin-releasing factor (CRF) (B) and vasopressin-associated neurophysin (NPII) in the macaque median eminence (ME). Immunoreactivity against copeptin (A; green) or CRF (B; green) and NPII (magenta) is merged in each right-hand *panel* (overlap; white). The outlined areas in merged images are enlarged. *Arrowheads* in (A) indicate tissue doubly immunopositive for copeptin and NPII. *Scale bars*, 50 μm, and 10 μm in enlarged images. InL, internal layer of the ME; ExL, external layer of the ME.

**Figure 10.**
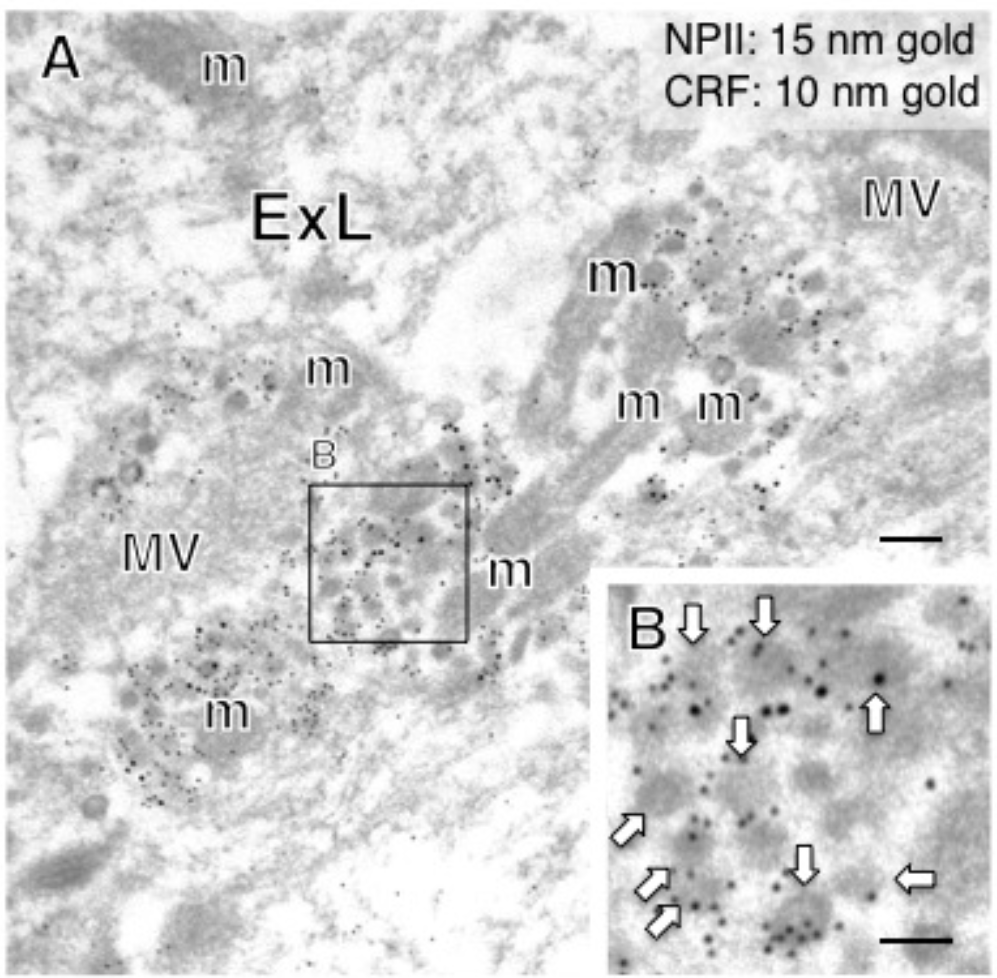
Double-label immunoelectron microscopy for vasopressin-associated neurophysin (NPII) and corticotrophin-releasing factor (CRF) in the external layer (ExL) of the macaque median eminence. In some axonal varicosities, numerous dense-cored neurosecretory vesicles were doubly immunopositive for CRF (10-nm gold particles) and NPII (15-nm gold particles). The outlined area in (A) is enlarged in (B). *Scale bars*, 200 nm, and 100 nm in the enlarged image. *Arrows* indicate dense-cored neurosecretory vesicles. m, mitochondrion; MV, clustered microvesicles.

**Figure 11.**
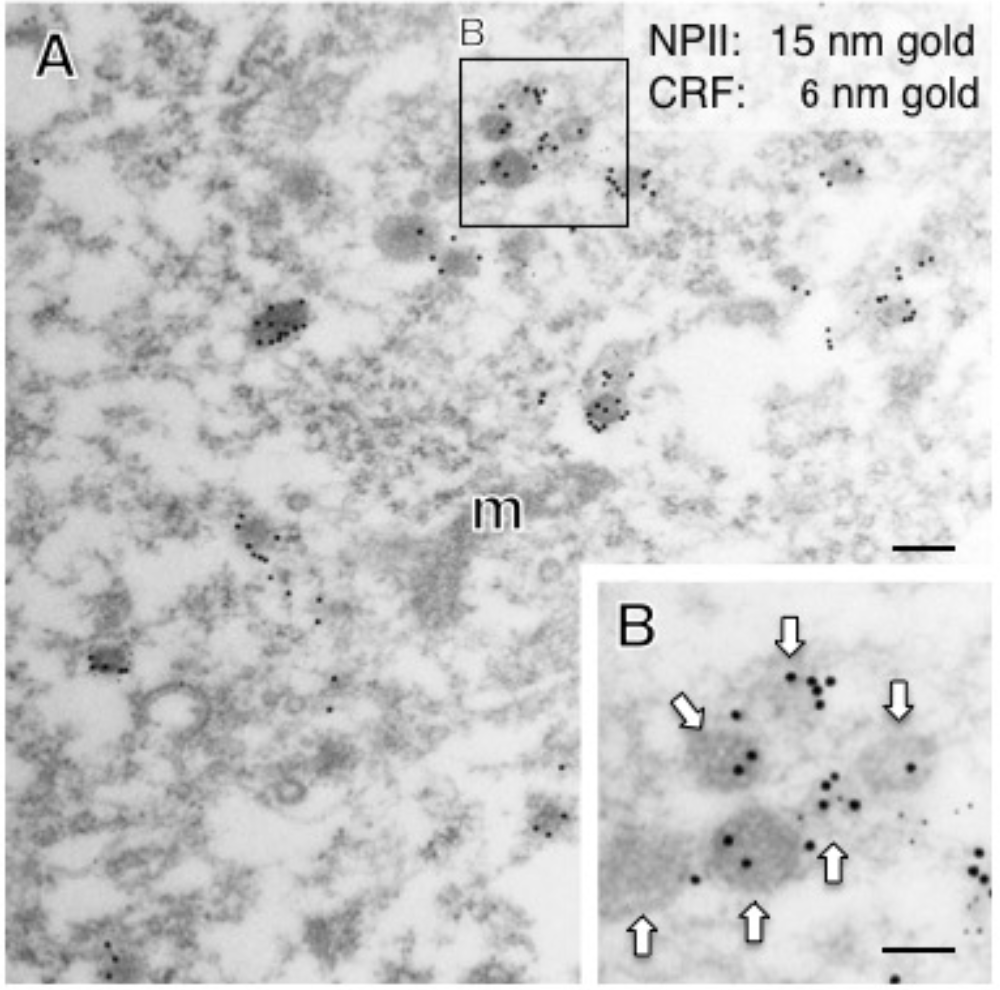
Double-label immunoelectron microscopy for vasopressin-associated neurophysin (NPII) and corticotropin-releasing factor (CRF) in the paraventricular nucleus (PVN) of the hypothalamus in a macaque monkey. (A) In the cell body of a possible parvocellular vasopressin neuron, many dense-cored neurosecretory vesicles located in the cell body were doubly immunopositive for CRF (6-nm gold particles) and NPII (15-nm gold particles). The outlined area in (A) is enlarged in (B). *Scale bars*, 200 nm, and 100 nm in the enlarged image. *Arrows* indicate dense-cored neurosecretory vesicles. m, mitochondrion.

### 2.5. Both magno- and parvocellular AVP neurons are glutamatergic

We first performed immunoelectron microscopy for L-glutamate in the posterior pituitary in order to determine its occurrence in the terminal regions of magnocellular AVP neurons. The immunoreactivity for glutamate was associated primarily with the membrane of small electron-lucent vesicles in the axonal endings of AVP neurons of posterior pituitary (Fig. 12A, B). In contrast, NPII-immunoreactivity was associated with dcv (Fig. 12A, B). Next, we examined the terminal regions of parvocellular axons in the median eminence. Triple staining for CRF/NPII/glutamate (Fig. 12C, D) showed that, as in the posterior pituitary, glutamate-immunoreactivity was associated with the membrane of electron-lucent microvesicles in the parvocellular endings which contained AVP/CRF-positive dcv (Fig. 12C, D) and in other similar endings of the terminals of other parvocellular neurosecretory neurons. Due to the mild formaldehyde fixation, the membrane of single microvesicles could not be clearly distinguished. However, a comparison with glutaraldehyde-fixed material leaves no doubt that these are clusters of microvesicles. Similar results were obtained by immunoelectron microscopy for VGLUT2 in both the posterior pituitary and the external layer of the median eminence (Fig. 13).

**Figure 12.**
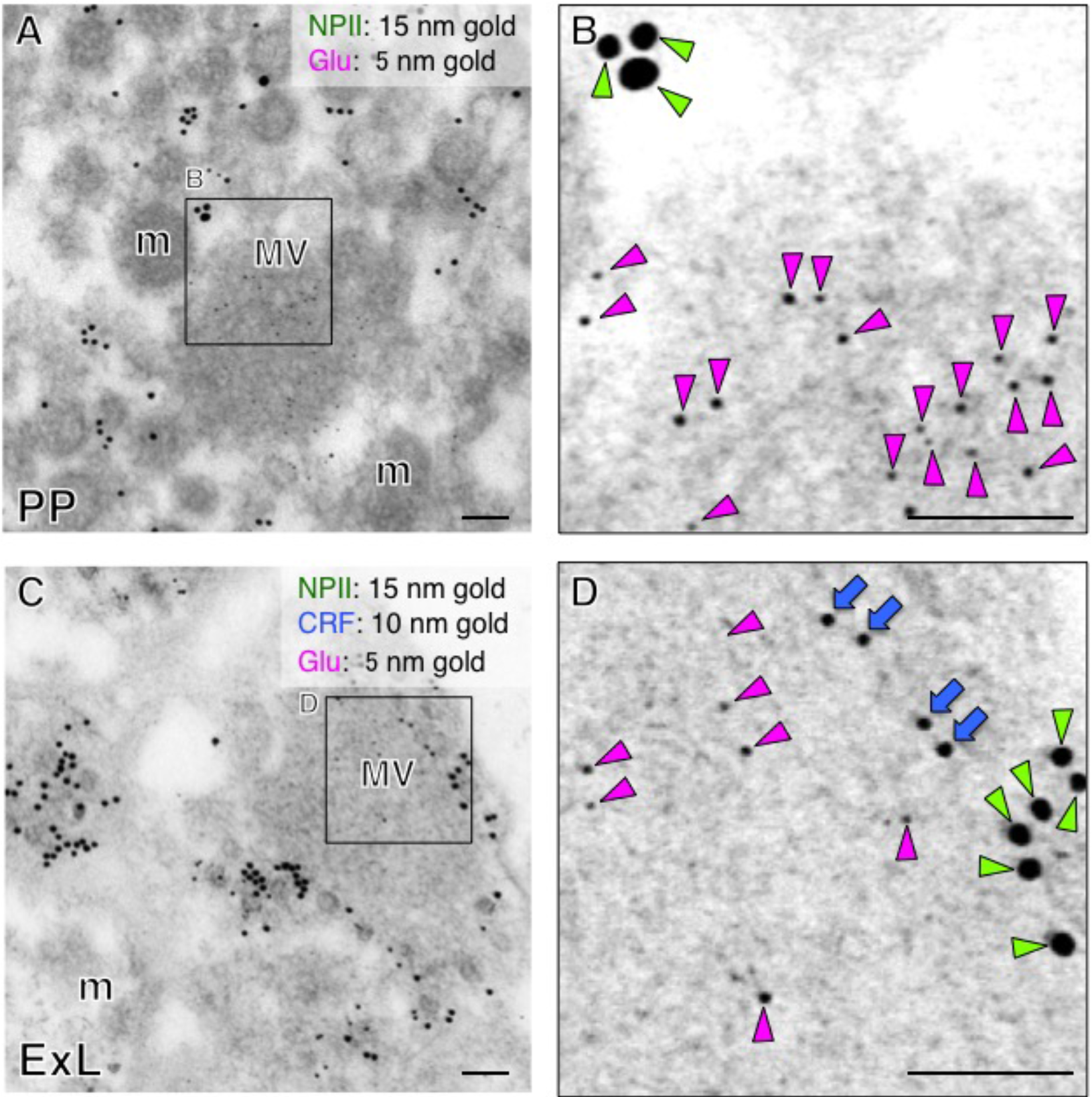
Immunoelectron microscopy for glutamate (Glu) in the posterior pituitary (PP) (A) and external layer of the median eminence (ExL) (C) in a macaque monkey. (B) Double-label immunoelectron microscopy for vasopressin-associated neurophysin (NPII) and Glu shows that Glu-immunoreactivity is associated with microvesicles in the macaque posterior pituitary. (D) Triple-label immunoelectron microscopy for corticotrophin-releasing factor (CRF)/NPII/Glu indicates that Glu-immunoreactivity is associated with electron-lucent small vesicles in the axonal endings, in the NPII/CRF-double positive endings. The outlined areas are enlarged. *Scale bars*, 100 nm. *Blue arrows* indicate NPII-immunoreactivity; *Green arrowheads* indicate CRF-immunoreactivity. *Magenta arrowheads* indicate Glu-immunoreactivity. m, mitochondrion; MV, clustered microvesicles.

**Figure 13.**
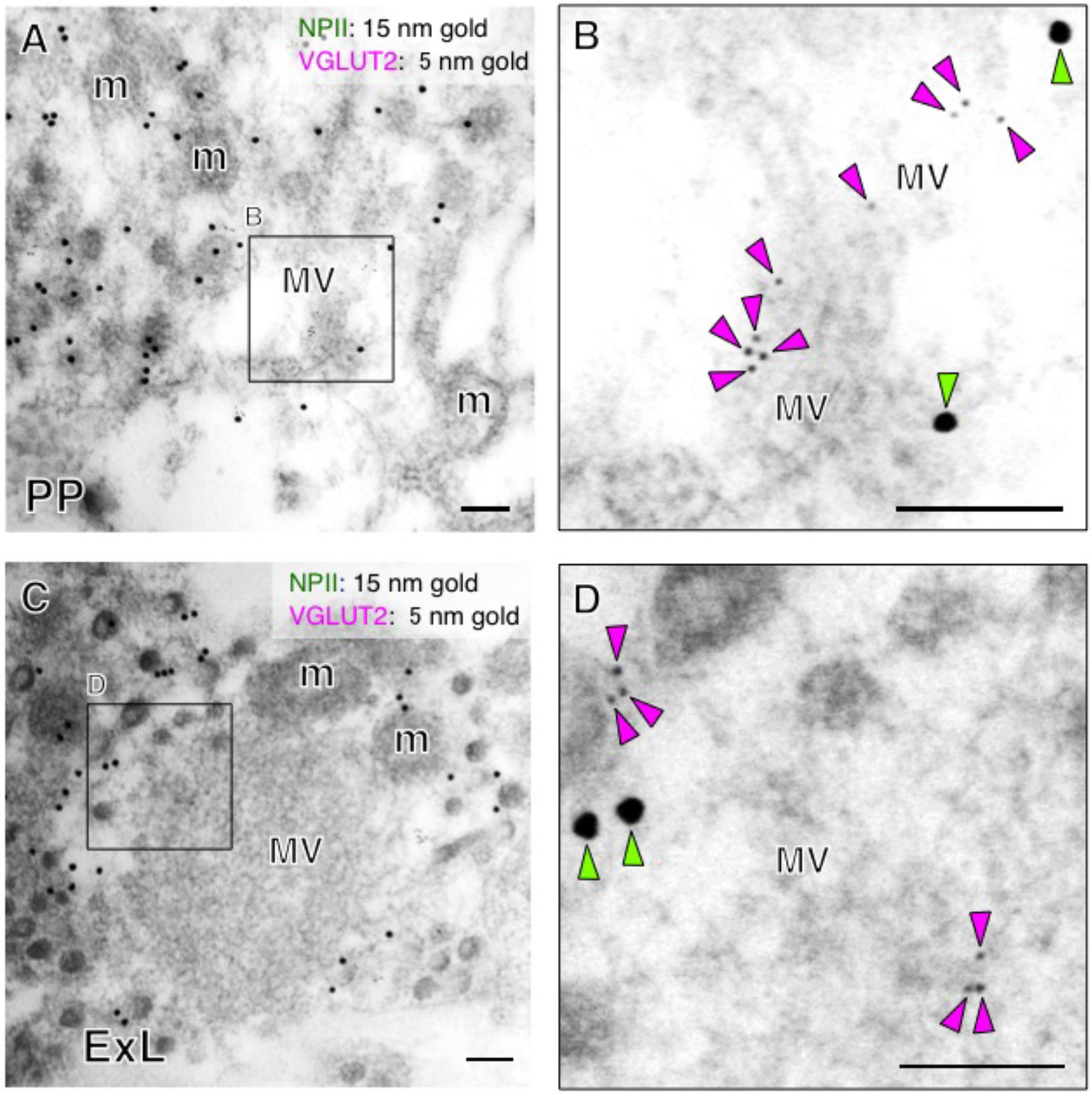
Immunoelectron microscopy for vesicular glutamate transporter 2 (VGLUT2) in the posterior pituitary (PP) (A) and external layer of the median eminence (ExL) (C) in a macaque monkey. (B) Double-label immunoelectron microscopy for vasopressin-associated neurophysin (NPII) and VGLUT2 shows that VGLUT2-immunoreactivity is associated with clustered microvesicles (MV) in the posterior pituitary. (D) Double-label immunoelectron microscopy for NPII/VGLUT2 indicates that VGLUT2-immunoreactivity is associated with the membrane of electron-lucent small vesicles in the vasopressin endings. The outlined areas are enlarged. *Scale bars*, 100 nm. *Green arrowheads* indicate NPII-immunoreactivity; *Magenta arrowheads* indicate VGLUT2-immunoreactivity. m, mitochondrion; MV, clustered microvesicles.

## 3. Discussion

It is often assumed that glutaraldehyde, a cross-linking fixative, is essential for the preservation of fine ultrastructure (Maunsbach, 1966a, 1966b) and that formaldehyde-only fixed tissues are inappropriate for electron microscopic analyses. However, the use of glutaraldehyde often results in a reduction of immunoreactivity and, for this reason, formaldehyde-fixed tissues without glutaraldehyde are used in most cases for immunohistochemical analysis at the light microscopic level (Maunsbach, 1966a, 1966b). The current study demonstrates that, in the macaque brain, interpretable ultrastructure and adequate ultrastructural immunoreactivity are both preserved in brain tissue even after long-term storage at –25°C for conventional light microscopy, provided that the fixed tissues are post-fixed with glutaraldehyde after the long-term storage. Thus, formalin-fixed macaque brains stored for a long period, at least 5-years, can be used for immunoelectron microscopic analysis, suggesting that not only macaque brain, but also postmortem formalin-fixed human brain tissue could be amenable to immunoelectron microscopic analysis. Furthermore, the ultrastructure of tissues from rare animals such as endangered species might be studied in this way even if they had been prepared for light microscopic immunohistochemistry and stored at –25°C in antifreeze solution.

The methodology used in this study has some obvious advantages in the ultrastructural analysis of the nervous system, in which tissue specimens are necessarily small, particularly where the brain is large, as in macaque monkeys and humans. After confirmation of the specificity of antibodies at the light microscopic level, post-embedding immunoelectron microscopic analysis can readily be performed on adjacent sections. Furthermore, because identification of brain nuclei and/or cells is easier at the light microscopic level, they can then be precisely targeted for post-embedding immunoelectron microscopy.

It has been reported that rodent AVP neurons also use glutamate as a neurotransmitter (Meeker et al., 1991; Meeker et al., 1989; Zhang et al., 2020; Ziegler et al., 2002). In this study, we examined the expression of glutamate and VGLUT2 in the macaque posterior pituitary. We first demonstrated by Western blotting that VGLUT2 was expressed in the macaque posterior pituitary, suggesting that magnocellular AVP (and also oxytocin) neurons use glutamate as a neurotransmitter in Japanese macaque monkeys. Immunoelectron microscopic analysis revealed that the AVP/CRF-immunoreactivity was restricted to the dcv, but the VGLUT2-immunoreactivity was associated with the smaller, electron-lucent microvesicles (Fig. 14). Similarly, immunoreactivity for glutamate itself was essentially restricted to microvesicles in the axonal endings of AVP neurons in both the posterior pituitary and external layer of the median eminence (Fig. 14). Because the antibody for glutamate, an amino acid, is raised against the cross-linking site of glutaraldehyde as the antigen, it is thought to be essential that glutaraldehyde is included in the fixative prior to glutamate immunocytochemistry (Meeker et al., 1991; Meeker et al., 1989; Sun et al., 2007). In our study, although glutaraldehyde was not included in the primary perfusion fixative, post-fixation of the formaldehyde-fixed tissue with glutaraldehyde enabled us to detect glutamate by immunoelectron microscopy even after prolonged storage. Therefore, the subcellular localization of amino acids appears to be preserved for a long-term even with conventional formaldehyde fixation. The immunoelectron microscopic detection of amino acids including glutamate and GABA should also be possible in conventional formaldehyde-fixed and stored human brain tissue.

**Figure 14.**
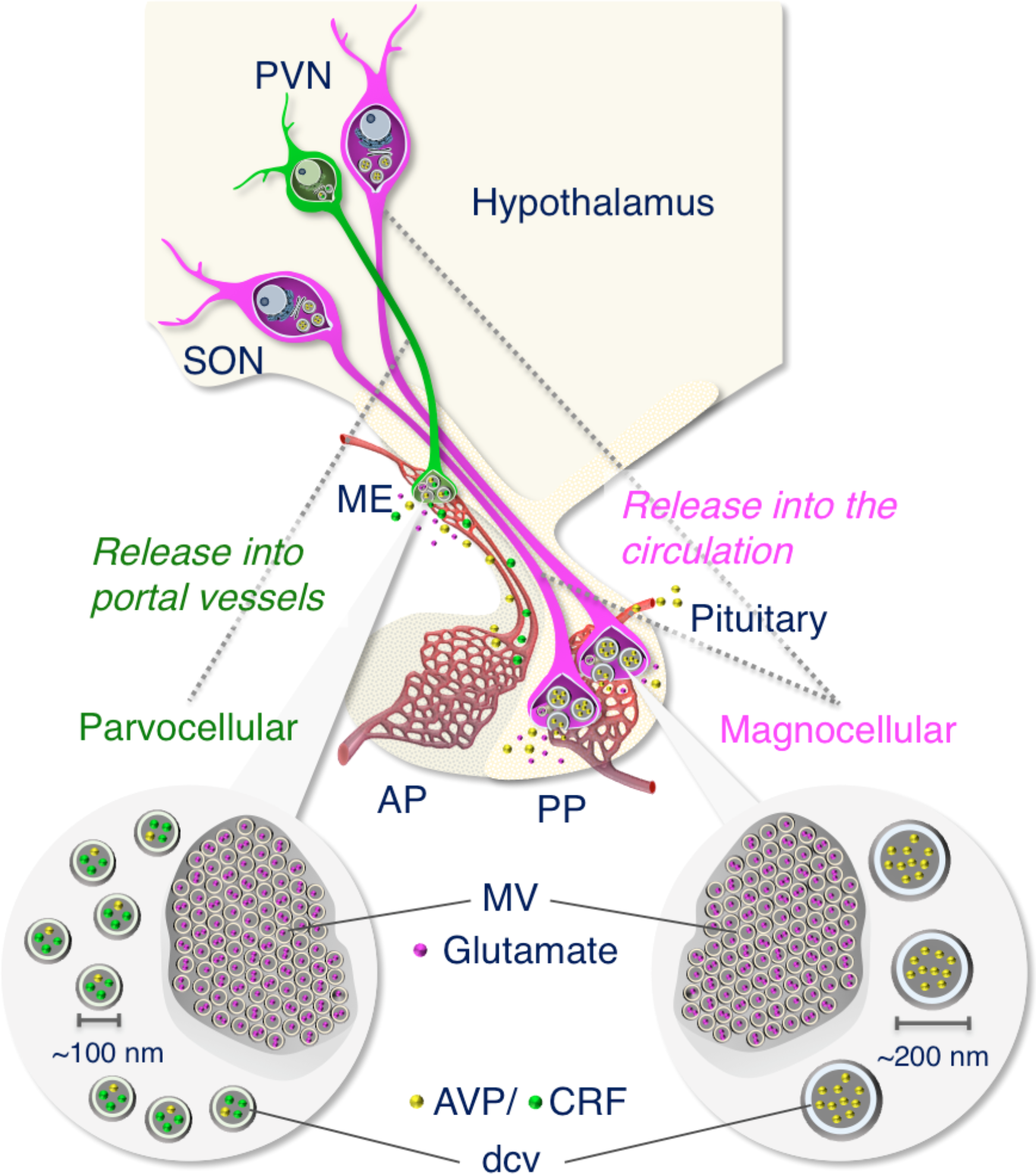
Working model showing that the size difference in dense-cored neurosecretry vesicles (dcv) between magnocellular (~200 nm) and parvocellular (~100 nm) vasopressin (AVP)- and/or corticotrophin-releasing factor (CRF)-producing neurons in the hypothalamus of the Japanese macaque monkey. We also demonstrate that both magno- and parvocellular AVP/CRF neurons are glutamatergic in primates. AP, anterior pituitary; ME, median eminence; MV, clustered microvesicles; PP, posterior pituitary; PVN, paraventricular nucleus; SON, supraoptic nucleus.

Our results in macaque tissue show that glutamate as an excitatory neurotransmitter is in microvesicles both in magno- and parvocellular AVP neurons. Although the localization of glutamate receptors has been reported in the neural lobe, median eminence and anterior pituitary, a clear role for glutamate secretion at these sites remains unknown (Brann, 1995; Hrabovszky & Liposits, 2007). Salt loading has been shown to produce robust increases in the VGLUT2 mRNA of AVP neuron perikarya and in VGLUT2-immunoreactivity in the rat posterior pituitary (Hrabovszky et al., 2006), suggesting that an osmotic challenge could produce an increase in glutamate release in the primate magnocellular AVP system (Fig. 14). Similarly, a role in the anterior pituitary of glutamate released from parvocellular AVP/CRF neurons into the hypothalamo-hypophysial portal veins remains unclear. Functional implications for glutamate release at the terminals of both magno- and parvocellular AVP neurons remain to be explored.

The size of dcv contained in the terminals of AVP neurons was quantified and compared among the posterior pituitary and internal/external layers of the median eminence. This showed, also for the macaque, the expected significant difference in size of the dcv (Fig. 14). It is known that the posterior pituitary and the internal layer of the median eminence contain the axons of magnocellular AVP neurons of the PVN and SON, whereas the external layer of the median eminence is the terminal region of parvocellular AVP neurons of the PVN (Buijs et al., 1983; Morris, 2020; Zimmerman et al., 1973) and other parvocellular neurons influence the anterior pituitary. In the dcv of the internal layer of the median eminence through which the magnocellular axons pass *en route* to the posterior pituitary, the AVP precursor should be processed sufficiently to reveal both prominent NPII- and copeptin-immunoreactivity in almost all of the axons (Kawata & Sano, 1982; Otubo et al., 2020; Zimmerman et al., 1973). In the macaque hypothalamus, quantifying only the size of the dcv in the cell body or its axon allows us to determine the magnocellular or parvocellular identify of the cell. Finally, our study confirms at the electron microscopic level that certain axonal endings in the external layer of the median eminence of macaques are derived from parvocellular AVP neurons of the PVN.

## 4. Conclusions

We report the immunoelectron microscopic characterization of AVP-producing neurons in primate hypothalamo-pituitary systems by the use of formaldehyde-fixed tissues stored at –25°C for several years. The size difference in dcv between magno- and parvocellular AVP neurons in Japanese macaque monkeys was determined (Fig. 14). We also demonstrate by immunoelectron microscopy that both magno- and parvocellular AVP neurons are glutamatergic in primates (Fig. 14). These results show, for the macaque brain, that interpretable neuronal ultrastructure and sufficient immunoreactivity for several relevant proteins are preserved even after a long-term storage of tissues fixed in formaldehyde for light microscopy. They indicate that this methodology could be applied to human post-mortem brain and, therefore, be very useful in translational research.

## 5. Materials and Methods

### 5.1. Animals

Three male (2–9-year-old, weight 2.3–12.6 kg) and four female (9–11-year-old, weight 7.2–10.8 kg) Japanese macaque monkeys (*Macaca fuscata*) were used in this study. Macaques were maintained in a temperature-controlled (22–24°C) room under a daily photoperiod of 12:12 hour light/dark cycle (lights off at 8:00 p.m.). These animals were checked and shown to be free of specific pathogens. Food and water were available *ad libitum*. All animals were kept in individual cages. The housing and experimental protocols followed the guidelines of the Ministry of Education, Culture, Sports, Science, and Technology (MEXT) of Japan, and were in accordance with the Guide for the Care and Use of Laboratory Animals prepared by Okayama University (Okayama, Japan), by Tokyo Medical and Dental University (Tokyo, Japan), and by National Institute for Physiological Sciences (Okazaki, Japan). All efforts were made to minimize animal suffering and reduce the number of animals used in this study.

### 5.2. Tissue processing

Hypothalamic sections and posterior pituitaries were obtained from four macaque monkeys (2 females and 2 males) as described previously (Otubo et al., 2020). Briefly, macaques were deeply anaesthetized with an overdose of sodium pentobarbital (~100 mg/kg body weight), and transcardially perfused with physiological saline followed by 4% paraformaldehyde (PFA) in 0.1 M phosphate buffer (PB, pH 7.4). After perfusion, brains and pituitaries were immediately removed and immersed in the same fixative for ~16 hours at 4°C. Coronal 30-μm-thick sections of the hypothalamus were prepared with a Linear-Slicer (PRO10, Dosaka EM, Kyoto, Japan). Preparations were rinsed with PB and then immersed in antifreeze solution (30% glycerol, 30% ethylene glycol, 0.36% NaCl, and 0.05% NaN_3_ in 40 mM PB) for storage at –25°C until use. For Western blot analysis, three female macaques were sacrificed by blood loss under deep pentobarbital anesthesia (see above). Posterior pituitaries were quickly removed, frozen immediately in powdered dry ice, and stored at – 80°C until use.

### 5.3. Immunofluorescence

The stored, formaldehyde-fixed sections were rinsed with phosphate-buffered saline (PBS) containing 0.3% Triton X-100 (PBST) five times for 10 minutes each. After blocking nonspecific binding with 1% normal goat serum and 1% BSA in PBST for 30 minutes at room temperature, the sections were incubated with mouse monoclonal anti-AVP-NPII antibody (PS41; 1:1,000 dilution) and with rabbit antiserum directed against copeptin (1: 10,000 dilution) or with rabbit antiserum directed against CRF (1: 20,000 dilution) for 4–5 days at 4°C. The mouse monoclonal NPII antibody has previously shown to be specific for AVP neurons in rodents (Castel et al., 1986; Wang et al.) and macaque monkeys (Otubo et al., 2020). The copeptin_7-14_ (ATQLDGPA) fragment, used as an antigen to produce our copeptin antiserum, is well conserved in mammals. It is identical in rats (Rehbein et al., 1986), mice (Hara et al., 1990), macaque monkeys (Otubo et al., 2020), and humans (Rehbein et al., 1986). The copeptin antiserum has also been characterized previously by our immunohistochemistry (Kawakami et al., 2021), Western (Satoh et al., 2015) and dot blot (Kawakami et al., 2021) analyses. The rabbit polyclonal antiserum against rat/human CRF (PBL rC70), which was kindly donated by Wylie Vale, has been characterized previously (Kawakami et al., 2021; Otubo et al., 2020; Sawchenko et al., 1984). It was directed against the rat form of CRF, which is identical to the human form, and has also been characterized in non-human primates (Foote & Cha, 1988; Otubo et al., 2020). Alexa Fluor 546-linked goat anti-mouse IgG (Molecular Probes, Eugene, OR, USA) and Alexa Fluor 488-linked goat anti-rabbit IgG (Molecular Probes) were used for detection at 1:1,000 dilution. Immunostained sections were viewed by confocal laser scanning microscopy (FluoView 1000, Olympus, Tokyo, Japan). The information for antibodies used in this study is shown in Table 1.

### 5.4. Post-embedding immunoelectron microscopy

The stored, formaldehyde-fixed sections were rinsed with PBS five times for 10 minutes each, and re-fixed in 4% PFA + 0.1% glutaraldehyde in 0.1 M PB for 4–6 hours at 4°C. Sections were then rinsed with 0.1 M PB three times for 5 minutes each, and were dehydrated through increasing concentrations of methanol, flat-embedded in LR Gold resin (Electron Microscopy Sciences, Hatfield, PA, USA) by gently pressing against the bottom of the flat-bottom-capsule with a pre-prepared resin block, and polymerized under UV lamps at −25°C for 24 hours. Ultrathin sections (70 nm in thickness) were collected on nickel grids coated with (or without for triple labeling) a collodion film, rinsed with PBS several times, then incubated with 2% normal goat serum and 2% BSA in 50 mM Tris(hydroxymethyl)-aminomethane-buffered saline (TBS; pH 8.2) for 20 minutes to block non-specific binding. The sections were then incubated with the mouse monoclonal antibody against NPII (1:200 dilution), with the rabbit polyclonal antisera against CRF (1:5,000 dilution), copeptin (1:100 dilution), L-glutamate (1:10 dilution, Abcam, Cambridge, UK), and/or with the guinea pig polyclonal antibody against VGLUT2 (1:20 dilution, Synaptic Systems, Göttingen, Germany) for 1 hour at room temperature. After incubation with the primary antibodies, the sections were washed with PBS, then incubated with a goat antibody against rabbit IgG conjugated to 5 or 10 nm gold particles (1:50 dilution, BBI Solutions, Cardiff, UK) and a goat antibody against mouse IgG conjugated to 15 nm gold particles (1:50 dilution, BBI Solutions) and/or a goat antibody against guinea pig IgG conjugated to 5 nm gold particles (1:50 dilution, Nanoprobes, Yaphank, NY, USA) for 1 hour at room temperature. The Abcam antibody to glutamate was raised in rabbit against L-glutamate conjugated to glutaraldehyde as the immunogen. The manufacturer’s information shows no measurable cross-reactivity detected against glutamate in peptides or proteins but modest cross-reactivity against D-glutamate. The specificity of this antibody was confirmed by immunostaining in the rat retina (Sun et al., 2007). To intensify the detectability of the immunoreaction for CRF in the cell body, a streptavidin-biotin intensification kit (Nichirei, Tokyo, Japan) was used. Tissues were first incubated with the biotinylated goat anti-rabbit IgG antibody for 10 minutes at room temperature, followed by incubation in avidin-biotin-horseradish peroxidase (HRP) complex solution for 5 minutes at room temperature. The sections were then washed with PBS, incubated with the goat antibody against HRP conjugated to 6 nm gold particles (1:50 dilution, Jackson ImmunoResearch Laboratory, West Grove, PA, USA) for 1 hour at room temperature.

Triple immunoelectron microscopy with antibodies against CRF, NPII, and glutamate was performed by using the front and back of ultrathin sections mounted on nickel grids without a supporting film. First, immunocytochemistry with a pair of primary antibodies (CRF and NPII) was performed on one side of the section and detected using 15 nm (mouse) and 10 nm (rabbit) colloidal gold particles (BBI Solutions), respectively. Next, immunocytochemistry with the other primary rabbit antibody (against glutamate) was performed on the other side of the section and detected by use of 5 nm colloidal gold particles (BBI Solutions).

Finally, the sections were contrasted with uranyl acetate and lead citrate and viewed using an H-7650 (Hitachi, Tokyo, Japan) or JEM-1010 (JEOL, Tokyo, Japan) electron microscope operated at 80 kV. The information for antibodies used in this study is shown in Table 2.

### 5.5. Western blotting

The lysates derived from posterior pituitaries were boiled in 10 μl sample buffer containing 62.5 mM trishydroxymethyl-aminomethane-HCl (Tris-HCl; pH. 6.8), 2% SDS, 25% glycerol, 10% 2-mercaptoethanol, and a small amount of bromophenol blue. These samples were run on a 4–20% SDS-PAGE and electroblotted onto a polyvinylidene difluoride (PVDF) membrane (Bio-Rad Laboratories, Hercules, CA, USA) from the gel by a semidry blotting apparatus (Bio-Rad Laboratories). The blotted membranes were blocked with PVDF Blocking Reagent for Can Get Signal (TOYOBO, Tokyo, Japan) for 30 minutes at room temperature and incubated overnight at 4°C with anti-AVP-NPII mouse monoclonal antibody (PS41) (1:1,000) or anti-VGLUT2 guinea pig polyclonal antibody (1:10,000) in Can Get Signal Solution 1 (TOYOBO). The blotted membranes were rinsed three times with 0.05% Tween 20 in Tris-HCl-buffered saline (TBST) and incubated with horseradish peroxidase-conjugated goat polyclonal antibody against mouse IgG (Bio-Rad Laboratories) or guinea pig IgG (Bioss Inc., Woburn, MA, USA) 1:10,000 dilution in Can Get Signal Solution 2 (TOYOBO) for 1 hour at room temperature. After washing for three times with TBST, blots were visualized by Immun-Star WesternC Chemiluminescence Kit (Bio-Rad Laboratories). The information for antibodies used in this study is shown in Table 2.

### 5.6. Statistical analysis

All data values were expressed as mean±standard error of the mean (SEM). Statistical significance was determined as *P* < 0.05 using one-way analysis of variance (ANOVA). When significant difference was found by ANOVA, the *post hoc* Tukey-Kramer test was performed. All the various analyses in this study were conducted “blind”.

## Author Contributions

Conceptualization, H.S.; methodology, A.O., S.M., T.O., K.S. and H.S.; validation, A.O. and H.S.; formal analysis, A.O. and H.S.; investigation, A.O., S.M., T.O., K.S. and H.S.; resources, T.O., K.S., Y.U. and H.S.; writing—original draft preparation, H.S.; writing—review and editing, J.F.M.; interpreted the data and provided advice and equipment, Y.U., J.F.M. and T.S.; supervision and project administration, H.S.; funding acquisition, T.O. and H.S. All authors have read and agreed to the published version of the manuscript.

## Acknowledgments

Tissues of Nihonzaru (Japanese macaque monkeys) were provided by National Institutes of Natural Sciences (NINS) and Kyoto University Primate Research Institute (KUPRI) with support in part by the National Bio-Resource Project (NBRP) of the Ministry of Education, Culture, Sports, Science and Technology (MEXT), Japan. We thank Dr. Narumi Katsuyama (Tokyo Medical and Dental University, Japan), Kei Tamaura and Ray Nomura for their technical support. We are grateful to Dr. Wylie Vale for the supply of an antiserum against CRF.

## Abbreviations

ANOVA: analysis of variance
AVP: arginine vasopressin
CRF: corticotrophin-releasing factor
dcv: dense-cored neurosecretory vesicle
HRP: horseradish peroxidase
NP: neurophysin
PB: phosphate buffer
PBS: phosphate-buffered saline
PFA: paraformaldehyde
PVDF: polyvinylidene difluoride
PVN: paraventricular nucleus
SEM: standard error of the mean
SON: supraoptic nucleus
Tris-HCl: trishydroxymethyl-aminomethane-HCl
TBST: Tris-HCl-buffered saline
VGLUT2: vesicular glutamate transporter 2

